# Contribution of CTCF binding to transcriptional activity at the *HOXA* locus in *NPM1*-mutant AML cells

**DOI:** 10.1101/2020.02.17.952390

**Authors:** Reza Ghasemi, Heidi Struthers, Elisabeth R. Wilson, David H. Spencer

## Abstract

Transcriptional regulation of the *HOXA* genes is thought to involve CTCF-mediated chromatin loops and the opposing actions of the COMPASS and Polycomb epigenetic complexes. We investigated the role of these mechanisms at the *HOXA* cluster in AML cells with the common NPM1c mutation, which express both *HOXA* and *HOXB* genes. CTCF binding at the *HOXA* locus is conserved across primary AML samples, regardless of *HOXA* gene expression, and defines a continuous chromatin domain marked by COMPASS-associated histone H3 trimethylation in *NPM1*-mutant primary AML samples. Profiling of the three-dimensional chromatin architecture of *NPM1*-mutant OCI-AML3 cells identified chromatin loops between the active *HOXA9*-*HOXA11* genes and loci in the *SNX10* gene and an intergenic region located 1.4Mbp upstream of the *HOXA* locus. Deletion of CTCF binding sites in OCI-AML3 cells reduced these interactions, but resulted in new, CTCF-independent loops with regions in the *SKAP2* gene that were marked by enhancer-associated histone modifications in primary AML samples. *HOXA* gene expression was maintained in the CTCF deletion mutants, indicating that transcriptional activity at the *HOXA* locus in *NPM1*-mutant AML cells does not require long-range CTCF-mediated chromatin interactions, and instead may be driven by intrinsic factors within the *HOXA* gene cluster.

## Introduction

The *HOX* genes encode developmentally regulated transcription factors that are highly expressed in acute myeloid leukemia (AML) and are important drivers of malignant self-renewal in this disease. Understanding how *HOX* genes are regulated in AML may therefore provide valuable information about the mechanisms that initiate and promote leukemogenesis. Previous studies have shown that expression of HOX family members in AML is nearly always restricted to specific genes in the *HOXA* and/or *HOXB* gene clusters (*HOXC* and *HOXD* genes are rarely expressed), and that expression patterns correlate with recurrent AML mutations (1). *HOX* expression is most closely associated with AMLs with *MLL* rearrangements, which exclusively express *HOXA* genes, and AMLs with the recurrent NPM1c mutation, which nearly always express both *HOXA* and *HOXB* cluster genes. The high prevalence of *NPM1* mutations make the combined *HOXA/HOXB* expression pattern the most common *HOX* phenotype in AML patients. However, the regulatory mechanisms that drive this expression pattern are poorly understood.

Studies of *HOX* gene regulation in model organisms have established that colinear expression of each *HOX* cluster is mediated by COMPASS/*Trithorax* and Polycomb group proteins (PcG), which promote gene activation and repression and perform methylation of histone H3 at lysine 4 (H3K4me3) and 27 (H3K27me3), respectively (2,3). These regulatory pathways are also involved in *HOX* gene regulation in AML cells, and are best understood for the *HOXA* cluster in AMLs with *MLL* rearrangements. MLL1 (KMT2A) is a component of the COMPASS complex, and MLL fusion proteins bind to the *HOXA* locus in AML cells and recruit the non-COMPASS histone H3 methyltransferase DOT1L, which is required for *HOXA* activation and AML development in *MLL*-rearranged leukemia models (4–7). Regulatory DNA elements that control three-dimensional chromatin architecture also play a role in *HOXA* gene regulation in AML cells. Specifically, the *HOXA* and *HOXB* clusters contain multiple binding sites for the chromatin organizing factor CTCF, and chromatin conformation experiments suggest these events mediate local chromatin loops in AML cells with *MLL* rearrangements (8). In addition, heterozygous deletion of a single CTCF binding site in the *HOXA* cluster in *MLL*-rearranged AML cells resulted in altered chromatin structure and reduced *HOXA* gene expression (9). These studies suggest that MLL fusion proteins directly activate the *HOXA* locus in ways that require specific CTCF binding events or their associated chromatin structures.

While these mechanistic insights have provided valuable information about *HOXA* regulation in *MLL*-rearranged AML, this molecular subtype accounts for <5% of all adult AML patients and only 25% of AMLs that express *HOXA* genes (1). Although AMLs with the *NPM1* mutations nearly always express *HOXA* genes (along with *HOXB* genes), it is unclear whether *HOXA* expression in these cells shares similar regulatory factors and chromatin structures that appear to be critical for *HOXA* expression in *MLL*-rearranged AML cells. In this study, we investigated histone modifications and chromatin domains at the *HOXA* locus in *NPM1*-mutated AML samples vs. other AML subtypes, and used a *NPM1*-mutant AML cell line model to determine whether CTCF binding at multiple sites at the *HOXA* locus is required to maintain *HOXA* gene expression and chromatin structure in *NPM1*-mutated AML cells.

## Materials and Methods

### Primary AML samples, normal hematopoietic cells, and AML cell lines

Primary AML samples and normal hematopoietic cells were obtained from bone marrow aspirates from AML patients at diagnosis or healthy volunteers following informed consent using protocols approved by the Human Research Protection Office at Washington University and have been described previously ((10,11); see Table S1). OCI-AML3 cells were cultured at 0.5-1 × 10^6^ cell/mL in MEM-alpha media with 20% FBS and 1% penicillin-streptomycin. The presence of NPM1c mutation in the parental OCI-AML3 line was verified by targeted sequencing at the beginning of the study and in RNA-seq data from wild type and mutant clones. Kasumi-1, IMS-MS2, and MOLM13 cell lines were cultured in RPMI-1640 with 1% penicillin-streptomycin and fetal calf serum (20% for Kasumi-1 and MOLM13, 10% for IMS-MS2).

### ChIP-seq preparation and analysis

ChIP-seq was performed using the ChIPmentation method (12) with the following ChIP-grade antibodies: CTCF (2899S), H3K27me3 (9733S), and H3K27ac (8173S) from Cell Signaling Technology and H3K4me3 (ab1012) from Abcam. All libraries were sequenced on a NovaSeq 6000 (Illumina, San Diego, CA) to obtain ∼50 million 150 bp paired end reads, and data were analyzed via adapter trimming with trim galore and alignment to the GRCh38 human reference sequence using bwa mem (13). Preparation of normalized coverage for visualization and analysis used the deeptools ‘bamCoverage’ tool (14), and peaks were called with macs2 using default parameters (15) for CTCF and epic2 (16) for all histone marks. Statistical comparisons were performed with DESeq2 (17) using raw fragment counts at identified peak summits. Visualizations were prepared using the Gviz package in R (18).

### Targeted deletion of CTCF binding sites via CRISPR-Cas9

Targeted deletions in OCI-AML3 cells were generated using CRISPR/Cas9 with guide RNAs from the UCSC genome browser (19,20) that overlapped CTCF ChIP-seq peaks from this study (see Table S2). Mutagenesis was performed via either an inducible Cas9-expressing OCI-AML3 cell line (Lenti-iCas9-neo vector; Addgene 85400) with lentiviral sgRNA expression (Addgene 70683), or transient transfection of OCI-AML3 cells with Cas9 protein complexed with synthetic tracrRNA/crRNA hybrids (Alt-R system, IDT, Coralville, IA). For the latter, RNAs were complexed with Cas9 protein using the manufacturer’s protocol with 1 million cells and 28 μM of Cas9/RNA for either transfection (CRISPRMAX; Thermofisher Scientific, Waltham, MA) or nucleofection (SG Amaxa Cell Line 4D-Nucleofector Kit, Lonza, Basel, Switzerland). Mutation efficiency was assessed in bulk cultures via DNA extraction, PCR with tailed primers (see Table S2), and sequencing to obtain 2×250 bp reads on an Illumina MiSeq instrument. Deletion abundance was assessed using custom scripts to analyze insertions/deletion in minimap2-based alignments (21). Single cell sorting into 96 well plates via FACS was used for expansion of individual mutant clones. Cells from single wells were screened via direct lysis by proteinase K (P8107S; NEB) in 20 μl of single cell lysis buffer (10 mM Tris-HCl pH 7.6, 50 mM NaCl, 6.25 mM MgCl2, 0.045% NP40, 0.45% Tween-20), PCR amplification, and gel electrophoresis; clones with evidence for deletions from the amplicon band size were sequenced, and clones with deletions were expanded for analysis.

### RNA analysis

RNA extractions were performed using ∼1 million cells using Quick-RNA MicroPrep Kit (Zymo Research, Irvine, CA). RT-qPCR for *HOXA9* used 100 ng of RNA for cDNA synthesis (Applied Biosystems, Foster City, CA) and qPCR in duplicate in a StepOnePlus PCR System (Applied Biosystems) for *HOXA9* exons 1-2 with a *GUSB* internal control (IDT). RNA-seq libraries were generated using 300 ng of RNA with the Kapa Hyper total stranded RNA library kit for Illumina (Roche) following the manufacturer’s instructions. Libraries were sequenced on a NovaSeq 6000 to obtain ∼50 million 2×150 bp reads. Reads were trimmed with trim galore, aligned with STAR (22), and transcript-level expression values in TPM were obtained with stringtie (23).

### In situ Hi-C

Hi-C libraries were prepared as described previously (24) using 4-5 million cells as input. Final libraries were first assessed by sequencing ∼1 million reads on a MiSeq instrument (using metrics recommended by Rao et al., (24)); passing libraries were sequenced to obtain >400 million 2×150 bp reads on a NovaSeq 6000. Hi-C data were analyzed on GRCh38 with juicer ((25); see Table S4). All analyses used contact matrices (mapping quality >30), deduplicated chimeric reads from the ‘merged_pairs.txt’ file, and chromatin loops and contact domains identified using HICCUPS and arrowhead, respectively, with default parameters (25). Chromatin loops from wild type and mutant OCI-AML3 cells were merged using bedtools ‘pairToPair’ function (26) with 5,000 bp overlap. Pairwise and joint comparisons of chromatin loop intensities were performed with hicCompare (27) and multiHicCompare (28), respectively, using raw contact counts from chromosome 7 obtained from the juicer hic file at 25 kbp resolution. Statistical comparisons used chromosome-wide FDR corrections for multiple hypothesis testing. Visualizations used the HiTC (29), GenomicInteractions and Gviz R packages.

### Data availability

All processed sequence data from this study (e.g., bigwig files for ChIP-seq, TPM values for RNA-seq, and Hi-C contact matrices) are available for public download at the following site: https://wustl.box.com/v/ghasemiCTCFHOXA. Raw sequence files for cell lines will be made available upon request.

## Results

### CTCF defines dynamic chromatin domains at the *HOXA* locus in primary AML cells

We previously showed that specific regions in the *HOXA* gene cluster have accessible chromatin in primary AML samples that coincide with CTCF binding sites in other human cell types (1). To confirm the presence of CTCF at these loci in primary AML cells, we performed CTCF ChIP-seq on bone marrow samples from AML patients with the NPM1c insertion mutation in *NPM1*, t(9;11) and t(11;19) *MLL* rearrangements, and t(8;21) creating the *RUNX1-RUNX1T1* gene fusion, which displayed the expected *HOXA* and *HOXB, HOXA* only, and no *HOX* expression patterns, respectively (Table S1, Figure S1A). This confirmed CTCF binding in chromatin-accessible regions between *HOXA6* and *HOXA7* (“CTCF Binding Site CBSA6/7”), and *HOXA7* and *HOXA9* (“CBSA7/9”), near the transcriptional start site of *HOXA10* (“CBSA10”), and in the 5’ UTR of *HOXA13* (“CBSA13”) (Figure 1A). Additional CTCF peaks with lower signals were present near the promoters of *HOXA4*, and *HOXA5*. All four major CTCF peaks were present in all AML samples regardless of *HOXA* expression status, and quantification of the CTCF signal demonstrated similar occupancy among the different AML types (Figures 1B-D).

**Figure 1.**
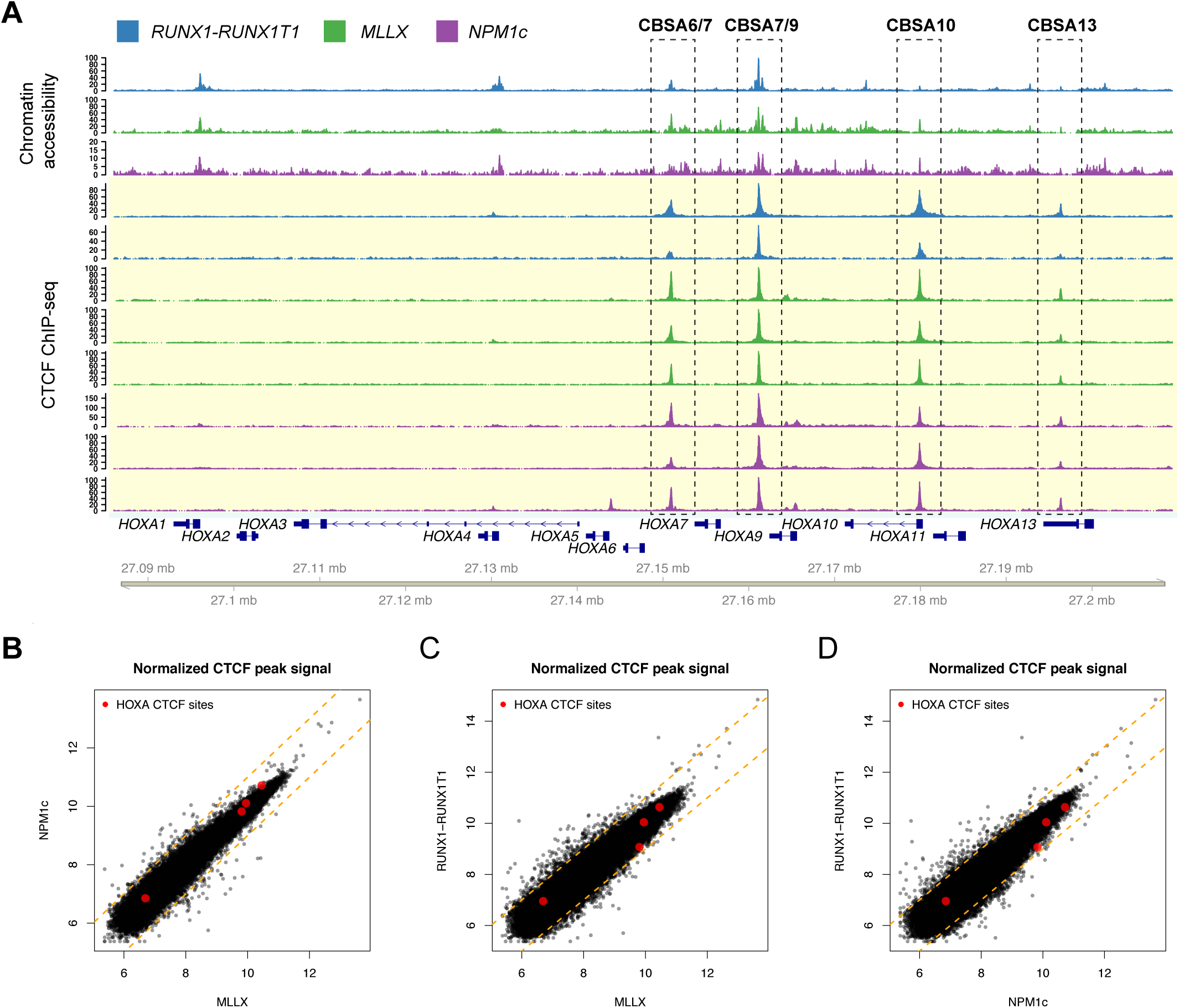
CTCF is bound to chromatin accessible sites at the *HOXA* locus in primary AML samples. A. Chromatin accessibility by ATAC (top tracks highlighted in white; ref. 1) and ChIP-seq for CTCF (bottom tracks highlighted in yellow) from primary AML samples with either t(8;21) creating the *RUNX1-RUNX1T1* gene fusion (blue), *MLL* rearrangements (green), or a normal karyotype and *NPM1* mutation (purple). CTCF sites CBSA6/7, CBSA7/9, CBSA10, and CBSA13 are indicated by the dashed boxes. B-D. Scatter plots comparing the CTCF peak summit counts between each AML type in log2 normalized read counts. Red points indicate ChIP-seq signal for the four CTCF binding sites highlighted in panel A, which are similar across AML samples. Dashed orange lines indicate a two-fold change between the samples.

We next determined whether these CTCF binding events defined chromatin domains in primary AML samples by performing ChIP-seq for H3K4me3 and H3K27me3 to measure active and repressed chromatin, respectively. This identified a distinct region of active chromatin in the center of the *HOXA* cluster that overlapped CBSA6/7 and CBSA7/9, and was conserved in the *MLL*-rearranged and *NPM1*-mutant AML samples, and normal CD34^+^ cells (which also express *HOXA* and *HOXB* genes) (Figures 2A, S2A). The H3K4me3 signal was continuous across this interval, including non-promoter sequences, and was also marked with H3K27ac in primary AML samples (Figure S2B). Regions adjacent to this active chromatin domain possessed repressive H3K27me3 marks in all AML samples and in CD34^+^ cells, which correlated with the expression levels of the overlapping genes (see Figures S1A, S2A). Histone ChIP-seq using AML samples with the *RUNX1-RUNX1T1* gene fusion and low *HOXA* expression showed that the active chromatin domain is dynamic, with little H3K4me3 signal and increased H3K27me3 in this AML type (Figure 2B; blue tracks). This was also observed in ChIP-seq data from FACS-purified normal promyelocytes and mature neutrophils (10) that also do not express *HOXA* genes (Figures 2B, pink and green tracks, Figure S2A). Interestingly, H3K4me3 was not completely absent in these cells, and similar H3K4me3 retention was observed in the *RUNX1-RUNX1T1* AML samples, although to a lesser extent. This suggests that regulation of *HOXA* genes in both AML cells and normal hematopoietic cells is associated with a poised chromatin state in CTCF-defined chromatin regions.

**Figure 2.**
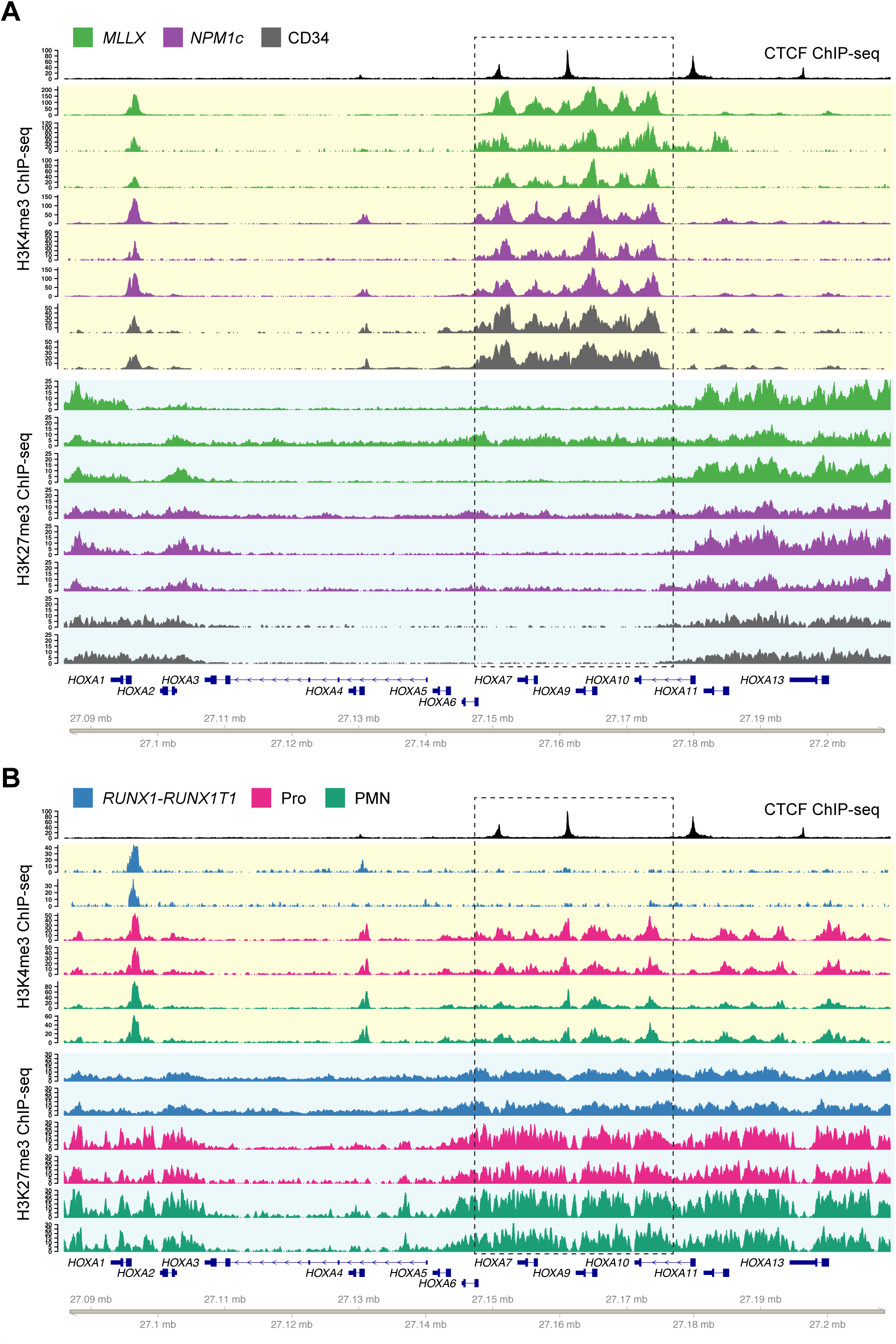
CTCF defines chromatin boundaries in AML samples and normal CD34^+^ cells with *HOXA* gene expression. A. Top track shows CTCF ChIP-seq from a *NPM1*-mutant primary AML sample. Tracks highlighted in yellow show ChIP-seq for H3K4me3 from primary samples with high *HOXA* expression, including AML samples with *MLL* rearrangements (green) and *NPM1* mutations (purple), and primary CD34^+^ hematopoietic stem/progenitor cells (HSPCs) purified from normal donor bone marrow samples (gray; GSE104579). Tracks highlighted in blue show H3K27me3 ChIP-seq from the same set of samples. B. H3K4me3 and H3K27me3 ChIP-seq from primary samples with no *HOXA* expression, including AML samples with t(8;21) creating the *RUNX1-RUNX1T1* fusion, and normal promyelocytes (CD14-, CD15+, CD16 low) and neutrophils (CD14-, CD15+, CD16 high) from healthy donor individuals. Dashed box indicates the region of dynamic chromatin that correlates with *HOXA* gene cluster expression.

### Targeted deletions at the *HOXA* locus eliminate CTCF binding but do not affect viability in *NPM1*-mutant OCI-AML3 cells

We sought to define the sequences that are required for CTCF binding at the *HOXA* locus, and determine whether loss of these DNA elements has functional consequences in *NPM1*-mutant AML cells. To this end, we used the OCI-AML3 cell line with a canonical *NPM1* insertion mutation and that expresses *MEIS1* and genes in both the *HOXA* and *HOXB* clusters (ref. 30; see Figure 3A). This pattern was unique to OCI-AML3 cells and was not observed in cell lines with other mutation-associated *HOX* expression phenotypes, including *MLL*-rearranged MOLM13 cells that expressed only *HOXA* genes, and the *RUNX1-RUNX1T1-*containing Kasumi-1 cell line, which had low *HOX* expression. ChIP-seq for CTCF using OCI-AML3 cells identified the four conserved CTCF sites observed in primary AML samples (Figures 3B, S3A) and peaks in the anterior *HOXA* cluster that were variably present in the primary AML samples (see Figure 1A). ChIP-seq for H3K4me3 and H3K27me3 also demonstrated an active chromatin domain between *HOXA9* and *HOXA13* (Figure 3B), which was consistent with the patterns of gene expression in this cell line, but different from chromatin domain boundaries in primary AML samples. However, the histone modifications between CBSA7/A9 and CBSA10 were shared between OCI-AML3 cells and *NPM1*-mutated primary AML samples, indicating that regulatory mechanisms in this specific region are conserved between primary AML samples and OCI-AML3 cells.

**Figure 3.**
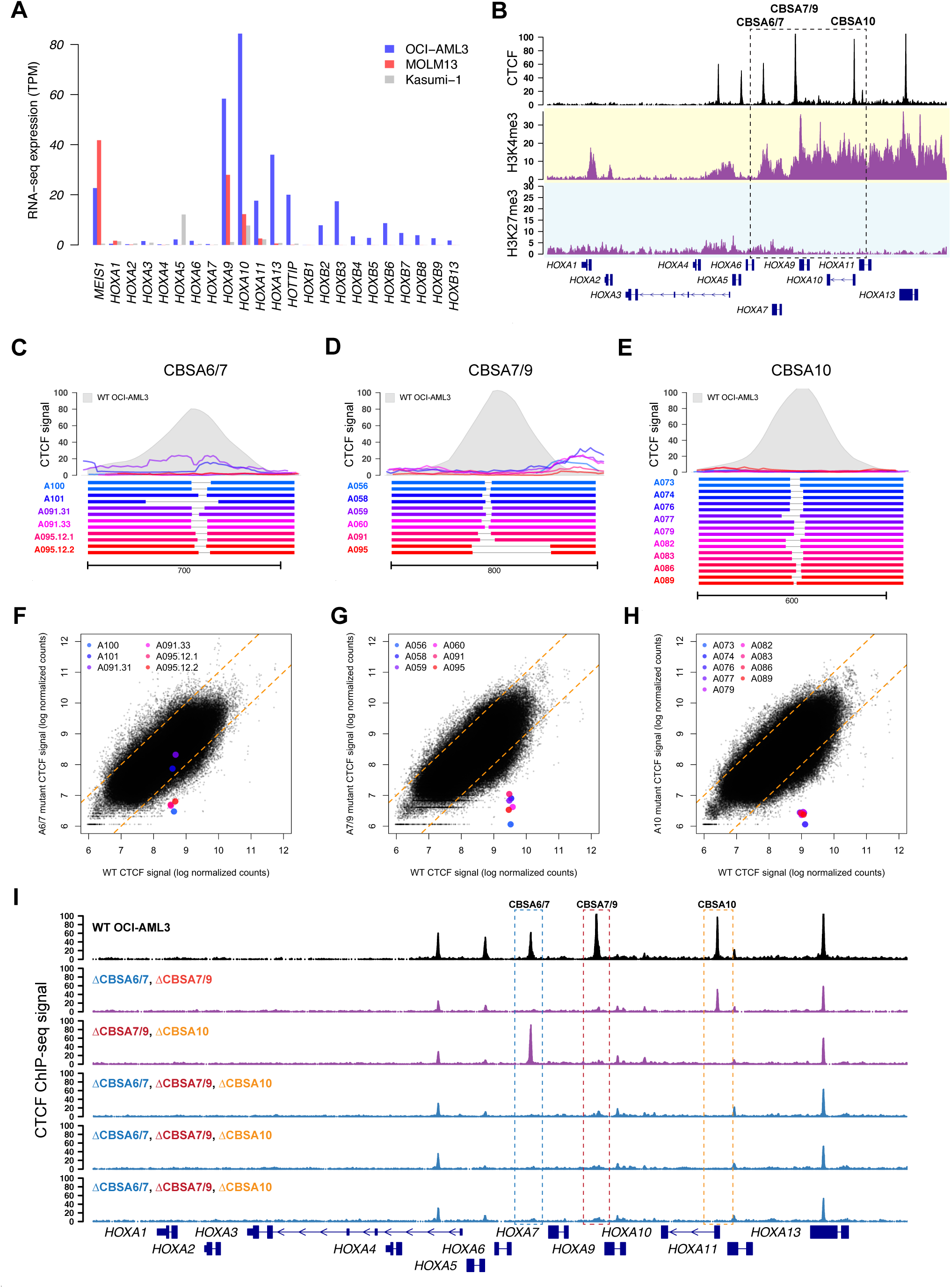
Targeted deletions eliminate CTCF binding in the *NPM1*-mutant OCI-AML3 cell line. A. RNA-seq expression of *HOXA* and *HOXB* genes in OCI-AML3 cells, which display the canonical mutant *NPM1*-associated *HOXA/HOXB* expression phenotype. Also shown are the *MLL*-rearranged MOLM13 cell line that expresses only *HOXA* genes, and the *RUNX1-RUNX1T1*-containing Kasumi-1 cell line that has low *HOXA* and *HOXB* gene expression. B. ChIP-seq data from OCI-AML3 cells for CTCF (black), H3K4me3 (highlighted in yellow) and H3K27me3 (highlighted in blue), which show conserved CTCF binding sites and distinct regions of active (H3K4me3) and repressed (H3K27me3) chromatin. C-E. Targeted deletions that disrupt CTCF binding in OCI-AML3 cells at sites CBSA6/7 (in C), CBSA7/9 (in D), and CBSA10 (in E). Bottom panels show allele pairs from homozygous or compound heterozygous deletion mutants at each site; top panels show CTCF ChIP-seq signal from these mutant cell lines (multi-colored lines) compared to wild type OCI-AML3 cells (in gray). F-H. Normalized CTCF ChIP-seq for all CTCF peaks from deletion mutants (Y axis) vs. wild type OCI-AML3 cells, showing dramatically reduced CTCF ChIP-seq signal in deletion mutants at all three sites, with the exception of clones A101 and A091.31, which only partially eliminates CTCF signal at site CBSA6/7. I. CTCF ChIP-seq tracks from double (in purple) and triple mutants (in blue), generated via sequential targeted deletion experiments. CTCF ChIP-seq from wild type OCI-AML3 cells is shown in black at the top for reference.

We next used CRISPR/Cas9-mediated editing to delete the three conserved CTCF binding sites in OCI-AML3 cells (CBSA6/7, CBSA7/9 and CBSA10). The resulting mutations did not appreciably alter markers of cell maturation in the edited cells (Figure S3B), and the deletion frequency at CBSA7/9 and CBSA10 remained stable after 7 and 14 days; CBSA6/7 deletion was less efficient, and showed a modest decrease in deletion frequency over time (Figure S3C). Cells sorted from bulk cultures were expanded and screened for deletions, which identified at least 5 individual clonal lines with homozygous deletions at each of the three sites (see Table S3), with deletions as small as 25, 10, and 9 bp sufficient to eliminate nearly all CTCF ChIP-seq signal from CBSA6/7, CBSA7/9, and CBSA10, respectively (Figure 3C-H). Additional experiments using single deletion mutants resulted in 10 doubly homozygous mutants and 7 triple mutants with homozygous deletions at all three sites (Figures 3I). None of the mutant OCI-AML3 clones showed overt defects in growth or maturation status, despite complete loss of CTCF binding in the posterior *HOXA* cluster.

### CTCF binding is not required for maintenance of *HOXA* gene expression or chromatin boundaries in *NPM1*-mutant AML cells

We selected 45 OCI-AML3 deletion mutants for *HOXA9* expression analysis via RT-qPCR to assess whether loss of CTCF binding affected *HOXA9* expression. Surprisingly, *HOXA9* expression was largely unchanged in all lines with deletions, with little evidence for consistent reduction in expression across the homozygous mutant clones and no trend toward decreased expression between wild type, heterozygous, and homozygous single mutants (Figure 4A) or multiply mutated clones (Figure 4B). Expression analysis via RNA-seq showed modest increases in the expression of the anterior *HOXA* genes *HOXA1*-*HOXA7* in lines with CBSA7/9 deletions, and subtle increases in the posterior *HOXA9-HOXA13* genes when CBSA10 was deleted (Figures 4E, 4D, S4A). However, most mutant lines showed few expression changes. Furthermore, a heterozygous single nucleotide polymorphism (SNP) in the *HOXA9* gene showed balanced expression of both alleles in wild type OCI-AML3 cells, and in all mutant lines, except for two with large mutations that involved the posterior *HOXA* cluster (Figure S4B).

**Figure 4.**
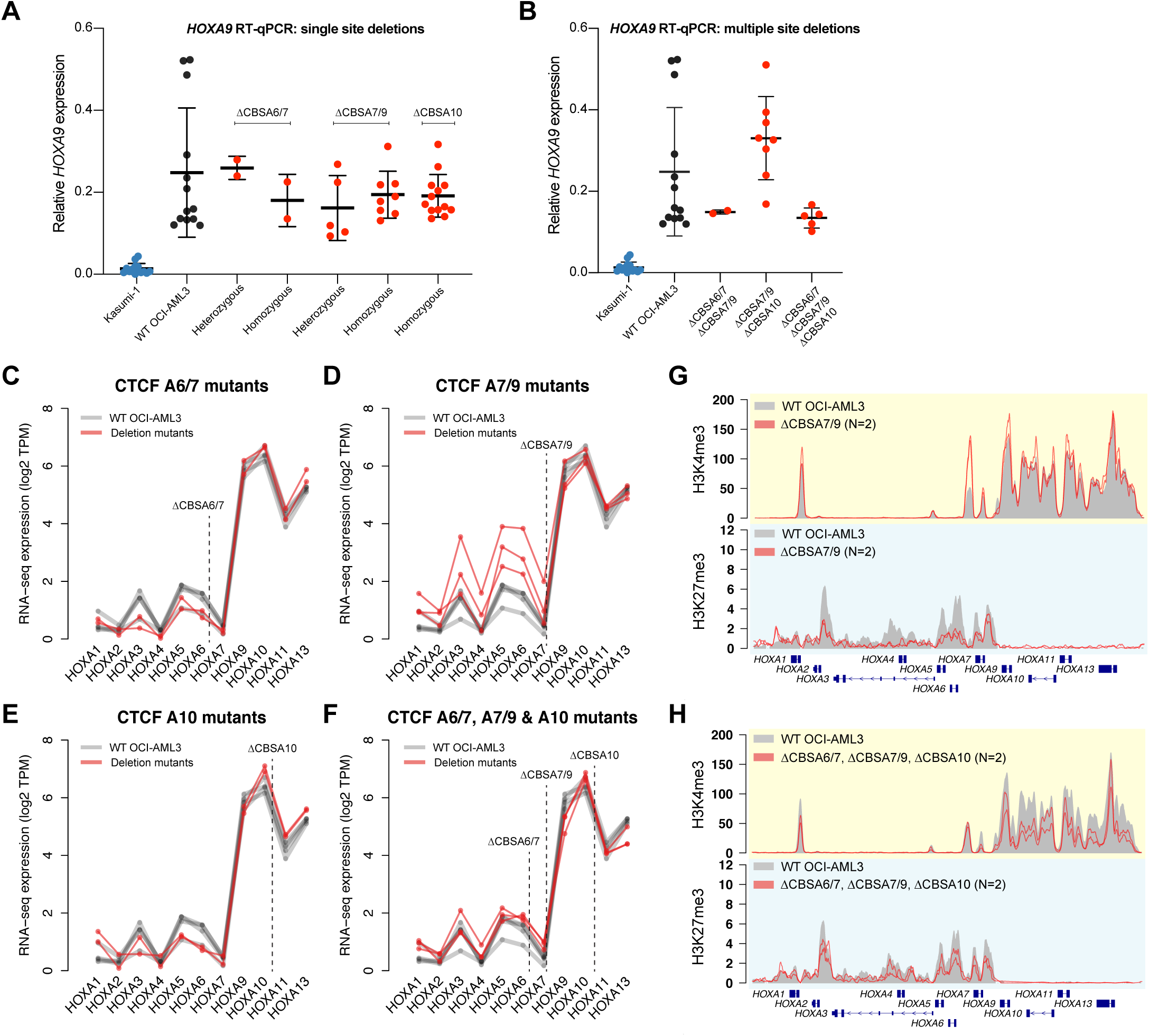
CTCF binding is not required to maintain gene expression or chromatin boundaries in the *HOXA* gene cluster. A. qPCR for *HOXA9* in single mutants with heterozygous or homozygous deletions at CBSA6/7, CBSA7/9, or CBSA10. *HOXA9* expression from the Kasumi-1 cell line is shown in blue as a no-*HOXA9* expressing control. B. qPCR for *HOXA9* in double and triple mutants at the CTCF binding sites indicated. Kasumi-1 cells are included in blue as in panel A. C-F. RNA-seq expression of all *HOXA* genes in single mutants (C-E) and triple mutants (D). Expression level is shown in log2 transcripts per million (TPM). G. ChIP-seq for H3K4me3 and H3K27me3 in deletion mutants lacking CTCF at site CBSA7/9 (in red; N=2). Mean ChIP-seq signal from wild type OCI-AML3 cells (N=2) is shown in gray. H. Mean ChIP-seq for H3K4me3 and H3K27me3 in triple mutants (N=2), with ChIP-seq from wild type cells shown in gray, as in panel G.

ChIP-seq for H3K4me3 and H3K27me3 was also performed on multiple mutants to determine whether loss of CTCF binding altered chromatin boundaries. H3K4me3 signal was reduced specifically at the CBSA7/9 site, but was otherwise intact. H3K27me3 was also still present, but was modestly decreased across the anterior *HOXA* genes in CBSA7/9 mutants; few changes were evident in other mutant lines, including triple mutants (Figure 4G, 4H, S4C-S4E). Mutants were also analyzed for other histone modifications, including H3K79me2 and H3K27ac, which were intact compared to wild type OCI-AML3 cells (Figure S4F).

### CTCF deletions result in compensatory *HOXA* chromatin loops

Given the role of CTCF in regulating chromatin architecture, we used *in situ* Hi-C (24) to define the chromatin interactions at the *HOXA* locus in wild type OCI-AML3 cells, CBSA7/9 single mutants, double mutants with deletions of CBSA6/7 and CBSA7/9 or CBSA7/9 and CBSA10, and two triply mutant lines (see Table S4). Analysis of the chromatin contacts from these cells identified a mean of 9,797 chromatin loops and 5,160 contact domains at 10 kbp resolution (ref. 25; Table S4). The *HOXA* cluster on chromosome 7p was located at the edge of a contact domain boundary (Figure 5A), with centromeric loops connecting *HOXA13* to the promoter of *TAX1BP1* and intronic sequences of *HIBADH* and *JAZF1*. The telomeric loops involved the remaining *HOXA* genes (*HOXA1*-*HOXA11*), which contacted regions in the *SKAP2*, and *SNX10* genes, and an intergenic region 1.4 Mbp upstream with no associated gene annotations. Similar chromatin loops were observed in Hi-C data generated from the *MLL*-rearranged MOLM13 cell line, and in previously reported data from normal human HSPCs (31) (Figures S5A, S5B), which both express *HOXA* genes.

**Figure 5.**
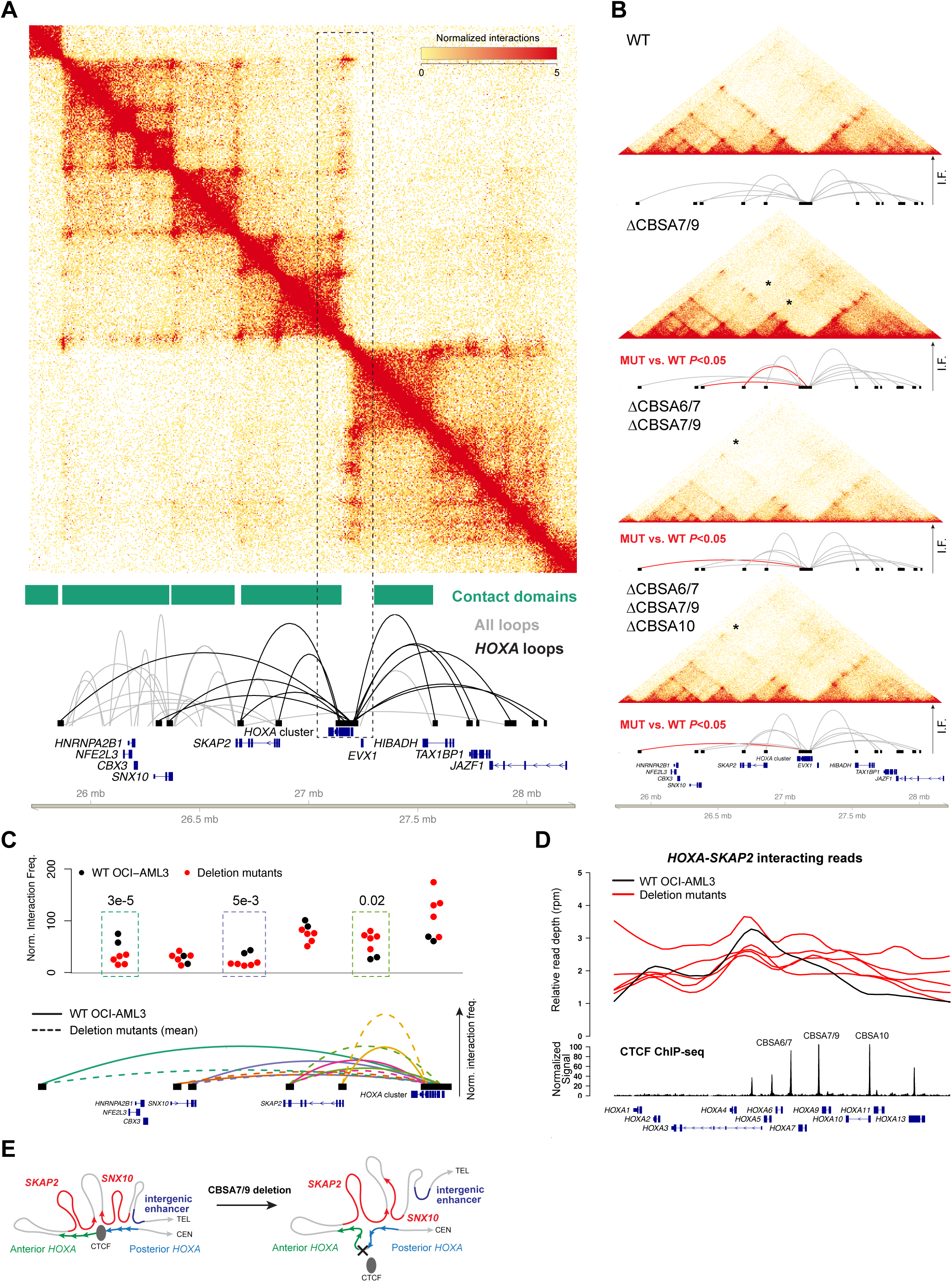
CTCF-mediated chromatin architecture at the *HOXA* locus in *NPM1*-mutant OCI-AML3 cells. A. Contact matrix, contact domains, and chromatin loops for a 2.5 Mbp region of chromosome 7p that contains the *HOXA* gene cluster from *in situ* Hi-C using wild type OCI-AML3 cells. Top panel shows the KR-normalized contact matrix at 5kb resolution. Tracks below the matrix show the contact domains and statistically supported chromatin loops identified using established methods (25). Loops shown in black are anchored within the *HOXA* gene cluster. B. Contacts and *HOXA* region loops from *in situ* Hi-C from wild type OCI-AML3 cells and homozygous (biallelic) single mutants at CBSA7/9, double mutants at CBSA6/7 and CBSA7/9, and triple mutants at CBSA6/7, CBSA7/9, and CBSA10. Loops in red indicate statistically significant differences in pairwise comparisons between each mutant and wild type cells with a chromosome-wide FDR<0.05. C. Normalized interaction frequencies for all datasets for 6 telomeric loops. Top panel shows normalized interaction frequencies for individual Hi-C libraries from wild type OCI-AML3 cells (N=2), single mutants at CBSA7/9, CBSA6/7-CBSA7/9 and CBSA7/9-CBSA10 double mutants, and two CBSA6/7-CBSA7/9-CBSA10 triple mutant lines (N=5 total mutant lines). Dashed boxes highlight loops that were statistically different in comparisons of all mutant libraries vs. two wild type OCI-AML3 libraries (chromosome-wide FDR<0.5). Mean interaction frequencies for chromatin loops from wild type and mutant lines are shown graphically in the lower panel in solid and dashed lines, respectively. D. Relative read depth (reads per million) across the *HOXA* locus for reads that interact between *HOXA* and *SKAP2*. Depth for wild type OCI-AML3 cells is shown in black and deletion mutants are shown in red. CTCF ChIP-seq signal from wild type OCI-AML3 cells and *HOXA* genes are shown in the bottom tracks. CTCF site deletions result in new interactions between the posterior *HOXA* cluster (genes *HOXA9*-*HOXA13*) with the *SKAP2* gene. E. Graphical representation of *HOXA* chromatin loops in wild type OCI-AML3 cells (left) and deletion mutants lacking *HOXA* CTCF binding sites (right).

We next analyzed Hi-C data from the OCI-AML3 deletion mutants to determine whether loss of CTCF binding altered chromatin architecture at the *HOXA* gene cluster. Although the general contact domain and chromatin loop structure remained intact, loops involving the *HOXA* locus were altered in the deletion mutants (Figure 5B). Pairwise comparisons of normalized chromosome 7 interaction frequencies between each mutant line vs. wild type cells identified decreased long-range interactions and increased interactions with the more proximal *SKAP2* gene (Figure 5B and S5C; red loops indicate chromosome-wide FDR<0.05). These findings were confirmed via statistical analysis of joint normalized contact frequencies from all samples, which demonstrated that mutant cells had significantly reduced interaction frequencies between the *HOXA* cluster and *SNX10* and the intergenic region, and increased interactions with two *SKAP2* loci (chromosome-wide FDR<0.05; Figure 5C). To define the specific *HOXA* genes involved in these changes, we mapped the positions of *HOXA-SKAP2* interacting reads within the *HOXA* locus. This showed that *SKAP2*-*HOXA* loops involved the anterior *HOXA* genes and intron 1 of *SKAP2* in wild type cells (Figure 5D, top panel, shown in black); however, in mutant cells there were increased interactions between the posterior *HOXA* genes *HOXA9-HOXA13* and intron 11 of *SKAP2* (Figure 5D, top panel, shown in red). Interacting reads with *SNX10* and the intergenic locus showed the opposite pattern, and were decreased in the posterior *HOXA* cluster (Figure S5D), indicating that loss of CTCF binding resulted in ‘spreading’ of the proximal *SKAP2* interactions to include the posterior *HOXA* genes (Figure 5E). There were no clear differences between cells with only CBSA7/9 deletions only, vs. double or triple deletions, suggesting that CBSA7/9 is critical for defining long-distance chromatin interactions with the anterior vs. posterior *HOXA* locus.

### Posterior *HOXA* chromatin loops correlate with expression and involve enhancer loci in primary AML samples

To determine the functional relevance of chromatin loops involving the posterior *HOXA* genes, we performed *in situ* Hi-C on the Kasumi-1 cell line with low expression of *HOXA* genes (see Figure 3A) to assess the correlation between chromatin architecture and posterior *HOXA* gene expression. Although the interactions in these cells displayed similar overall patterns in the *HOXA* region, the posterior *HOXA* genes did not participate in any statistically supported chromatin loops (Figures 6A, 6B), and reads involved in *HOXA* contacts mapped primarily to the anterior *HOXA* genes (Figure 6C). Loops to the posterior *HOXA* cluster were also present in the *NPM1*-mutant IMS-M2 cell line with high *HOXA* expression (Figures S6A, S6B), further supporting the observation that expression of these *HOXA* genes is associated with long-distance chromatin interactions in *NPM1*-mutant AML cells.

**Figure 6.**
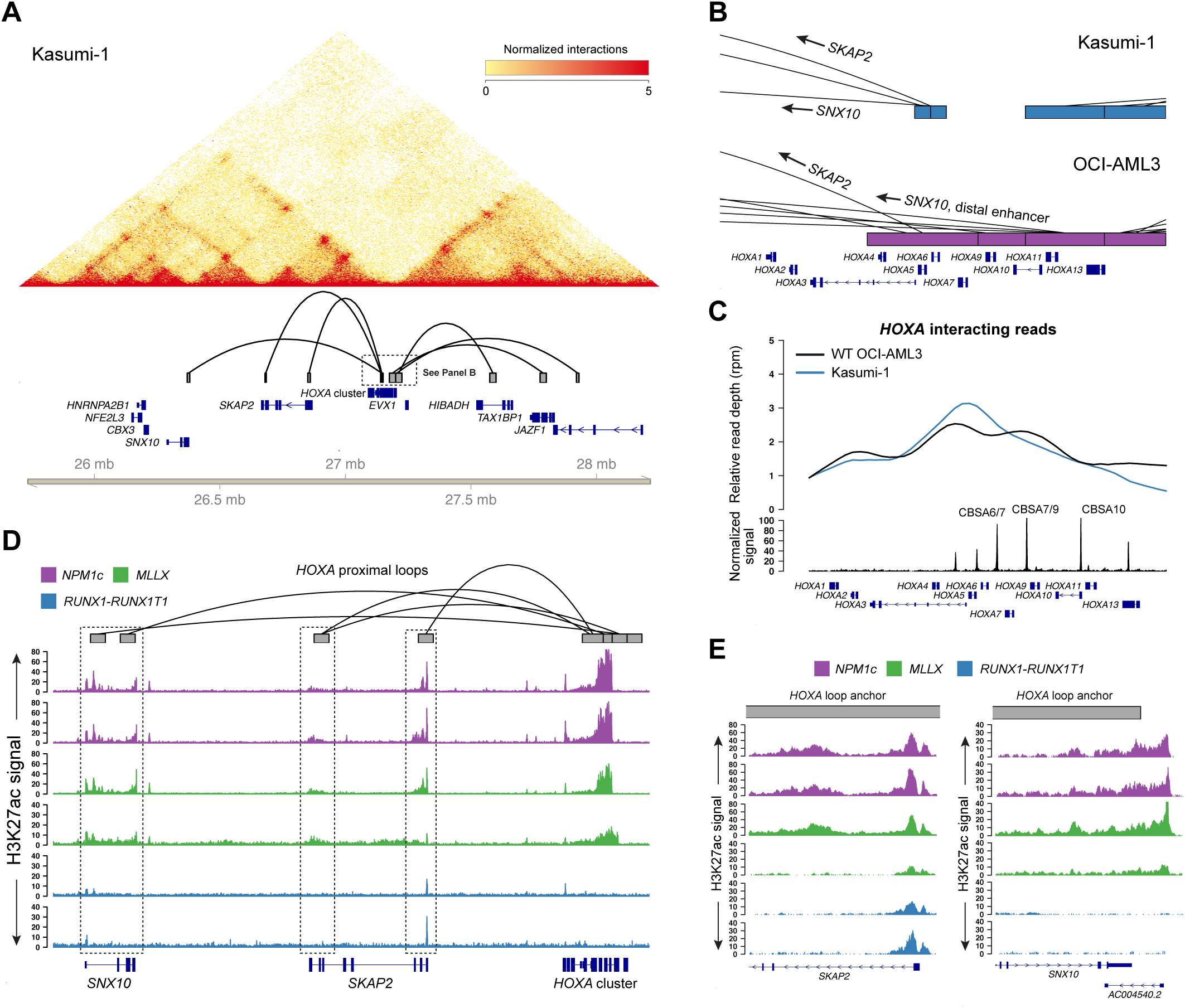
Long-distance interactions with posterior *HOXA* genes correlate with expression and involve loci with enhancer-associated histone acetylation. A. Chromatin interactions at chromosome 7p from the Kasumi-1 AML cell line that contains t(8;21)/*RUNX1*-*RUNX1T1* and has low expression of *HOXA* genes. The *HOXA* cluster resides at the boundary of a topologically associated domain, but genes in the posterior *HOXA* cluster do not participate in chromatin loops in this cell line. B. Focused view of *HOXA* chromatin loops in the Kasumi-1 and OCI-AML3 cell lines with low and high posterior *HOXA* expression, respectively, which display the differences in chromatin loop structure and loop anchors. C. Relative read depth in reads per million of Hi-C reads between the *HOXA* locus and telomeric chromatin loops in Kasumi-1 (in blue) and OCI-AML3 cells (in black). Interacting reads in Kasumi-1 cells are localized to the central *HOXA* cluster, compared to interacting reads in OCI-AML3 cells that map to the central and posterior *HOXA* locus. D. Enhancer-associated histone H3 lysine 27 acetylation (H3K27ac) at *HOXA* interacting loci from primary AML samples with and without *HOXA* gene expression. Primary AML samples include patients with NPM1c (purple; N=2 distinct patients) and *MLL* rearrangements (t(9;11) and t(11;19)) (green; N=2 distinct patients) with high *HOXA* expression, and samples with t(8;21)/*RUNX1*-*RUNX1T1* (N=2 distinct patients). Regions highlighted in the dashed boxes were shown to interact with the *HOXA* cluster in the OCI-AML3 cell line model (loop track, top), and display H3K27ac signal suggesting they may represent functional genomic elements. E. High-resolution view of two loop anchor regions in *SKAP2* intron 1 and downstream of the *SNX10* gene, which possess the enhancer-associated H3K27ac mark in *HOXA*-expressing AML samples but not in samples with no *HOXA* expression.

We next analyzed ChIP-seq data for H3K27ac from primary AML samples to determine whether loci that interact with *HOXA* genes may have enhancer properties. Indeed, both intron 1 and 11 of *SKAP2* displayed H3K27ac signals in *NPM1*-mutant AML samples, as well as *MLL*-rearranged samples that also express *HOXA* genes (Figure 6D). Multiple loci in the *SNX10* gene also possessed this mark, including a non-coding RNA downstream of the 3’ UTR of *SNX10* (Figure 6E), but H3K27ac was not present at other regions that formed contacts with the *HOXA* cluster, including the distal intergenic locus (Figure S6C). Although the enhancer modifications observed at interacting regions had relatively low signal, they were clearly absent in the *RUNX1-RUNX1T1* samples, which lack *HOXA* expression. This provides evidence that these specific regions interact with the *HOXA* cluster in primary AML samples that express *HOXA* genes, and may therefore contribute to *HOXA* gene regulation.

## Discussion

*HOX* transcription factors are drivers of self-renewal in AML cells and are highly expressed in AMLs with *NPM1c* mutations. In this study, we demonstrated that the chromatin organizing factor CTCF is bound equally to specific sites at the *HOXA* locus in *NPM1*-mutant primary AML samples compared to other AML subtypes, and define a dynamic chromatin domain in primary AML samples and normal hematopoietic cells. Targeted deletions in the *NPM1*-mutant OCI-AML3 cell line eliminated CTCF binding, but surprisingly did not disrupt *HOXA* gene expression. This was true when multiple binding sites were deleted individually, or in combination, and was supported by independent mutant clones that showed little evidence for consistent decreases in *HOXA* gene expression or changes to the histone modifications we measured. However, loss of CTCF binding did result in clear alterations to the chromatin loops involving the posterior *HOXA* genes, including *HOXA9* and *HOXA10*. Specifically, long-range loops were diminished, and were replaced by compensatory interactions with regions of the *SKAP2* gene. Some of these candidate enhancers have been reported in other studies (32), but have not previously been shown to be active in hematopoietic cells. Further investigation of these sequences, and their associated regulatory proteins, may shed light into factors that promote *HOXA* gene activation in normal and malignant myeloid cells.

The central observation in this study is that CTCF binding at the *HOXA* cluster is not absolutely required for maintenance of *HOXA* expression in *NPM1*-mutated AML cells. Deletion of CTCF site CBSA7/9 was previously shown to affect *HOXA* expression in the *MLL*-rearranged MOLM13 cell line (9). These discrepant findings may be due to fundamental differences in how *HOX* genes are regulated in *MLL*-rearranged vs. *NPM1*-mutant AML cells. Indeed, the *HOX* expression phenotype of these AML subtypes is strikingly different: *MLL* rearrangements are associated with only *HOXA* gene expression, whereas both *HOXA* and *HOXB* genes are expressed in *NPM1*-mutant AML cells. *HOXA* and *HOXB* genes are simultaneously downregulated during normal myeloid maturation (1), which implies that the clusters may be controlled by common factors that may function in different ways in *NPM1*-mutant vs. *MLL*-rearranged AML cells. We also observed differences in *HOXA* cluster expression patterns in MOLM13 vs. OCI-AML3 cells, including higher expression of *HOXA11, HOXA13*, and the *HOTTIP* long noncoding RNA in OCI-AML3 cells. *HOTTIP* has an established role in *HOXA* gene regulation (33,34), and its higher expression may influence the requirement for certain CTCF-mediated chromatin architectures, thereby making these CTCF binding sites dispensable for steady-state expression of posterior *HOXA* genes in *NPM1*-mutant AML cells.

Three-dimensional architecture is an important component of the regulatory control of *HOXA* genes during development. Our analysis of Hi-C data from OCI-AML3 cells identified long-range loops between the active posterior *HOXA* genes *HOXA9*-*HOXA13* and sequences with enhancer-associated epigenetic marks in the *SNX10* and *SKAP2* genes. Additional loops were identified at other loci, but these did not possess active chromatin modifications in primary AML samples. AML cells without *HOXA* expression did not display these loops, and lacked active histone marks at the interacting loci, which provides evidence that these contacts may be functionally relevant for *HOXA* gene expression in AML. Interestingly, loss of CTCF within the *HOXA* cluster revealed the plasticity of these interactions, implying that multiple loci may serve as exchangeable *HOXA* enhancers, which is reminiscent of other well-studied, complex regulatory systems, and has been implicated in developmental *HOXA* gene regulation (35,36).

Our data showed that transcriptional activity and active histone modifications remained highly localized to the posterior *HOXA* cluster in OCI-AML3 cells, even after elimination of key CTCF boundaries. In fact, there was a tendency for deletion mutants to show increased *HOXA* gene expression levels compared to wild type cells. This suggests the posterior *HOXA* cluster possesses intrinsic properties that maintain its active state. It is unclear whether this activity is caused by the compensatory interactions we observed, or if these chromatin loops are consequences of persistent gene expression that is mediated by autonomous, sequence-specific factors that recruit transcriptional machinery. The *HOXA* cluster contains many highly conserved noncoding elements beyond CTCF binding sites. Additional studies will be required to better understand whether these elements provide key signals that promote and maintain *HOXA* gene expression in *NPM1*-mutant AML cells.

## Acknowledgements

This work was supported by the American Society of Hematology (ASH Scholar Award), the American Cancer Society (Institutional Research Grant), and the National Cancer Institute (K08CA190815) to D.H.S. Primary AML samples were provided by the Genomics of AML Program Project (P01CA101937, T. Ley, PI). Technical assistance was provided by the Siteman Cancer Center Tissue Procurement and Cell Sorting cores (NCI Cancer Center Support Grant P30CA91842).

