## Supplemental Figures for "Contribution of CTCF binding to transcriptional activity at the *HOXA* locus in *NPM1*-mutant AML cells"

**Supplemental Figure Legends**

**Figure S1. *HOXA* and *HOXB* gene cluster expression in primary AML samples.** Heatmap showing RNA-seq expression of the *HOXA* and *HOXB* cluster genes from primary AML samples with t(8;21) creating the *RUNX1-RUNX1T1* fusion, t(9;11) or t(11;19) with *MLL* rearrangements, and the NPM1c mutation in *NPM1* in log2 transcripts per million (TPM) from ref. 1. These recurrent mutations are associated with canonical HOX expression patterns, including little to no *HOX* expression in t(8;21) samples, *HOXA* genes only in *MLL*-rearranged samples, and combined *HOXA/HOXB* expression in *NPM1*-mutant samples.

**Figure S2. *HOX* gene expression and histone modifications in primary AML samples and normal hematopoietic cells.** A. *HOXA* and *HOXB* expression in purified bone marrow cells from normal donors, including CD34+ hematopoietic stem/progenitor cells (HSPCs), promyelocytes, and neutrophils (1). *HOXA* and *HOXB* genes are expressed in HSPCs and then downregulated in promyelocytes and neutrophils. B. ChIP-seq signal for H3K27ac at the *HOXA* locus from a subset of the primary AML samples shown in Figure S1, including samples with t(8;21), t(9;11) or t(11;19), and the NPM1c mutation. High H3K27ac signal is present at the active *HOXA* chromatin domain shown in Figure 2.

**Figure S3.** A. CTCF ChIP-seq in the *NPM1*-mutant OCI-AML3 cells compared to primary AML samples with *NPM1* mutations. Top track (black) shows normalized CTCF ChIP-seq signal from OCI-AML3 cells. Bottom tracks (purple) show data from three distinct primary AML samples. Red dashed box highlights three CTCF binding sites that are conserved between the OCI-AML3 cell line and primary AML samples. B. Representative flow cytometry for CD11b and CD14 from wild type OCI-AML3 cells and a triple mutant with deletions at CTCF binding sites CBSA6/7, CBSA7/9, and CBSA10. C. Mean percentage of CD11b, CD14 double positive cells in wild type

OCI-AML3 cells (N=4) and mutant clones with single deletions (CBSA6/7, N=4; CBSA7/9, N=5; CBSA10, N=5), double deletions (CBSA6/7+CBSA7/9, N=9; CBSA7/9+CBSA10, N=13), and triple deletion mutants (N=9), which demonstrates little difference in these flow markers in the mutant OCI-AML3 clones. Bars show mean percent double positive cells +/- one standard deviation. D. Deletion allele fraction over time in bulk edited cultures where CBSA6/7, CBSA7/9, or CBSA10 were targeted with CRISPR/Cas9. DNA was prepared from edited cultures on days 2, 7, and 14 for PCR amplification and direct sequencing of amplicons. Reads were mapped to the genome and counted if they were unmodified or contained deletions >10 bp. Bars represent the fraction of reads with deletions and error bars show the 95% confidence interval of the point estimate.

**Figure S4.** A. Heatmap representation *HOXA* cluster genes from RNA-sequencing of wild type OCI-AML3 cells (N=4) and mutant clones with homozygous deletions of CBSA6/7, CBSA7/9, or CBSA10, and double mutants (CBSA6/7+CBSA7/9, N=1; CBSA7/9+CBSA10, N=2), and triple mutants (N=3). Expression values are indicated in absolute normalized read counts obtained from the 'vst' function in DESeq2. B. Allelic ratio of a heterozygous common SNP in wild type OCI-AML3 cells and mutant clones from RNA-seq data, demonstrating that expression of both alleles is balanced in nearly all clones with deletions, including a mutant with a heterozygous deletion, and multiple compound heterozygotes. Note that double mutants A91-10 (*) and A95-5 (**) show skewed expression because these clones contain a 29 kbp deletion and a 29 kbp inversion, respectively, that disrupt CTCF binding sites but also involve the *HOXA9* gene. C. ChIP-seq for H3K4me3 (yellow panel) and H3K27me3 (blue panel) from a mutant OCI-AML3 clone with deletion of CBSA6/7. Mean ChIP-seq signals from 2 replicates of wild type OCI-AML3 cells are shown in gray. D. Histone ChIP-seq for a mutant clone with deletion of CBSA10, displayed as C. Histone ChIP-seq from double mutant clones with deletion of CBSA7/9 and either CBSA6/7 (N=1) or CBSA10 (N=2). F. ChIP-seq for histone H3K79 dimethylation

(H3K79me2; top panel in orange) and H3K27 acetylation (bottom panel in white). Data are from three mutant clones with deletions of either CBSA7/9 (N=2) or CBSA10 (N=1) (shown in blue). Data from wild type OCI-AML3 cells are shown in gray.

**Figure S5.** A. Chromatin contact matrix for chromosome 7p for the *MLL*-rearranged MOLM13 cell line from *in situ* Hi-C data. Data were generated using the approach described in the Methods section for comparison with Hi-C data from OCI-AML3 cells. Chromatin interactions involving the *HOXA* gene cluster are similar between MOLM13 and OCI-AML3 cells. Data shown were normalized using KR normalization and were visualized using juicebox (see ref. 25). B. Chromatin contact matrix for chromosome 7p from normal hematopoietic stem/progenitors (HSPCs) from reference 31. Data are presented as in A. C. Chromatin contact matrix and comparative loop analysis for a double mutant OCI-AML3 clone lacking CTCF binding sites CBSA7/9 and CBSA10 (see Table S3). The loop shown in red was found to be significantly different from wild type OCI-AML3 cells in a pairwise comparison of normalized interaction frequencies (see Methods). D. Normalized read depth of interacting reads between the *HOXA* cluster and the *SNX10* gene and the distal intergenic locus from wild type (black) and mutant (red) OCI-AML3 cells. Note that these interactions involve the posterior *HOXA* cluster in wild type cells, but in mutant clones lacking CTCF binding sites these interactions are reduced and occur across the *HOXA* cluster.

**Figure S6.** A. Chromatin contact matrix for chromosome 7p from the *NPM1*-mutant IMS-MS2 cell line. B. Focused view of chromatin loop anchors at the *HOXA* locus in IMS-MS2 cells compared to OCI-AML3 cells. Note the loop anchors involve the posterior *HOXA* cluster in both cell lines, compared to Kasumi-1 cells which do not express *HOXA* genes (See Figure 6B). C. ChIP-seq for H3K27ac from primary AML samples with NPM1c (purple), *MLL* rearrangements (N=1 each of t(9;11) and t(11;19); green), and t(8;21) and the *RUNX1-RUNX1T1* gene fusion for a 480 kbp region that includes *HOXA* interacting regions in the *SNX10* gene and the distal intergenic locus, which is demarcated by the dashed box. Note that the intergenic locus does not possess the enhancer associated H3K27ac modification, and therefore does not appear to be functionally active in primary AML samples that express the *HOXA* genes.

Figure S1

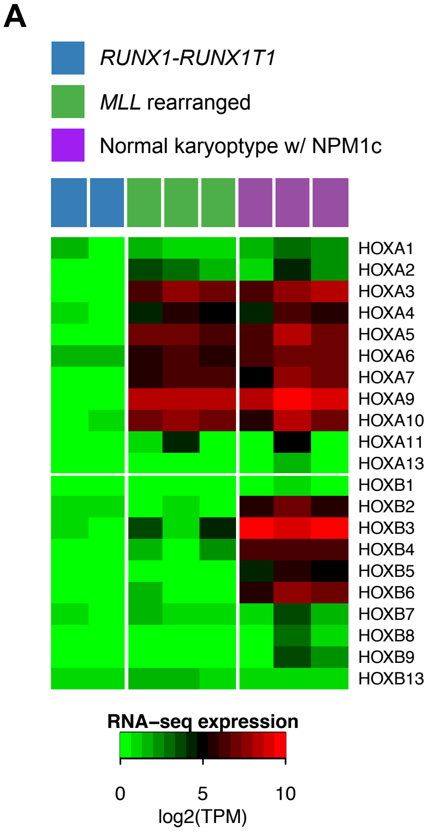

A
RUNX1-RUNX1T1
MLL rearranged
Normal karyoptype w/ NPM1c
HOXA1
HOXA2
HOXA3
HOXA4
HOXA5
HOXA6
HOXA7
HOXA9
HOXA10
HOXA11
HOXA13
HOXB1
HOXB2
HOXB3
HOXB4
HOXB5
HOXB6
HOXB7
HOXB8
HOXB9
HOXB13
RNA–seq expression
0
5
10
log2(TPM)

### Figure S2

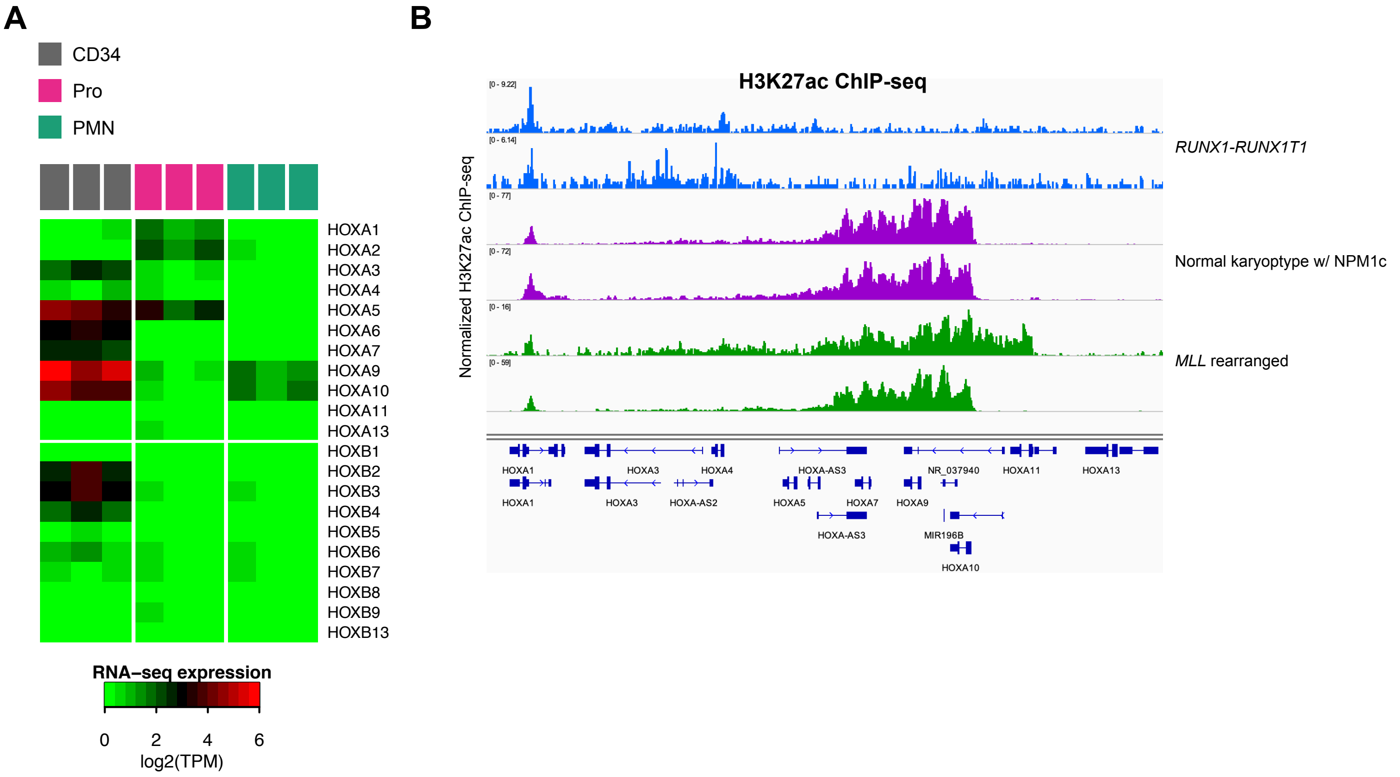
A
CD34
Pro
PMN
HOXA1
HOXA2
HOXA3
HOXA4
HOXA5
HOXA6
HOXA7
HOXA9
HOXA10
HOXA11
HOXA13
HOXB1
HOXB2
HOXB3
HOXB4
HOXB5
HOXB6
HOXB7
HOXB8
HOXB9
HOXB13
RNA–seq expression
0
2
4
6
log2(TPM)
B
H3K27ac ChIP-seq
Normalized H3K27ac ChIP-seq
RUNX1-RUNX1T1
Normal karyoptype w/ NPM1c
MLL rearranged
HOXA1
HOXA3
HOXA4
HOXA-AS3
NR_037940
HOXA11
HOXA13
HOXA1
HOXA3
HOXA-AS2
HOXA5
HOXA7
HOXA9
HOXA-AS3
MIR196B
HOXA10

**Figure S3**

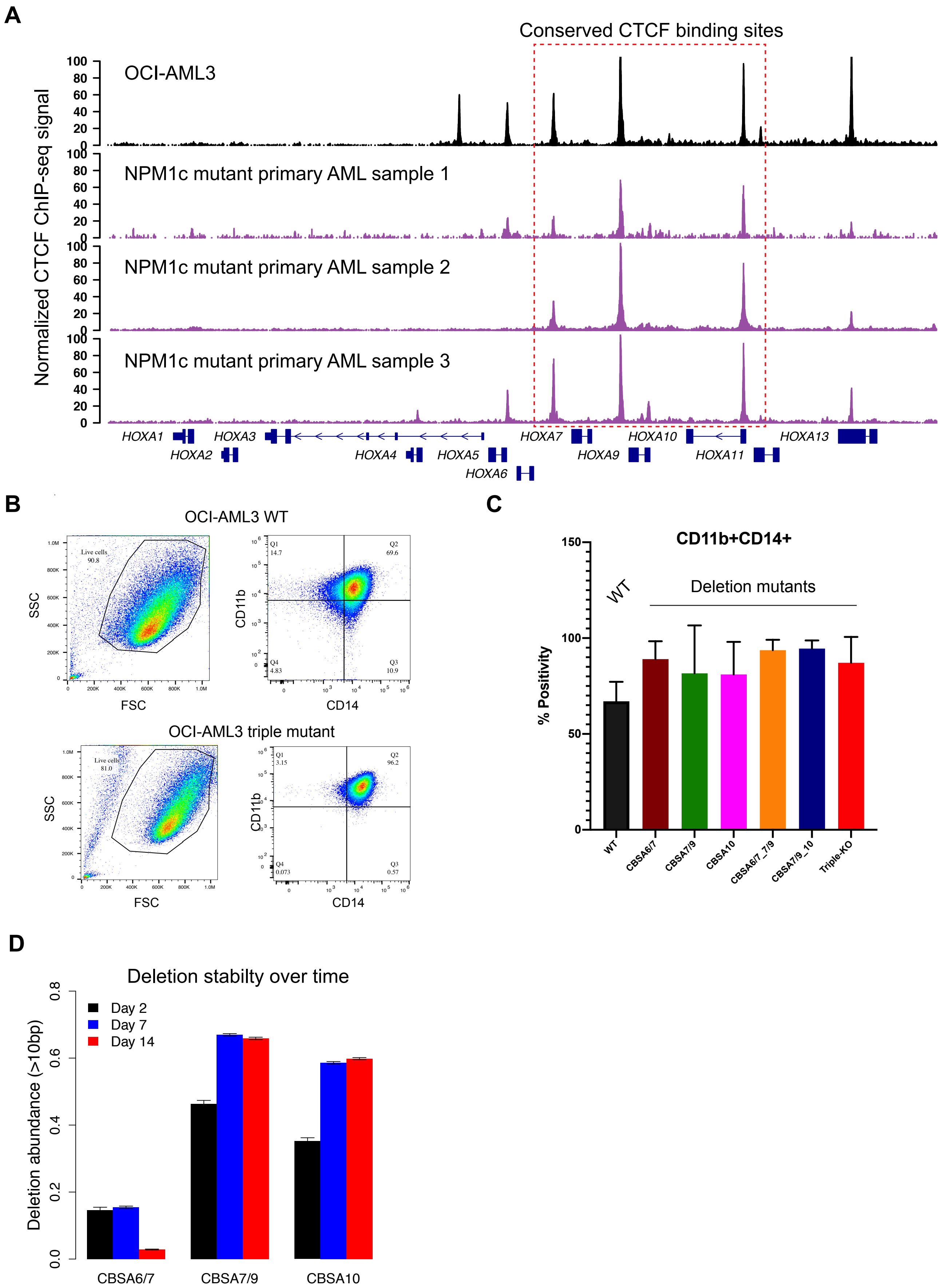
A
Conserved CTCF binding sites
Normalized CTCF ChIP-seq signal
OCI-AML3
NPM1c mutant primary AML sample 1
NPM1c mutant primary AML sample 2
NPM1c mutant primary AML sample 3
HOXA1
HOXA2
HOXA3
HOXA4
HOXA5
HOXA6
HOXA7
HOXA9
HOXA10
HOXA11
HOXA13
B
OCI-AML3 WT
SSC
FSC
CD11b
CD14
Live cells 90.8
Q1 14.7
Q2 69.6
Q3 10.9
Q4 4.83
OCI-AML3 triple mutant
Live cells 81.0
Q1 3.15
Q2 96.2
Q3 0.57
Q4 0.073
C
CD11b+CD14+
% Positivity
WT
Deletion mutants
WT
CBSA6/7
CBSA7/9
CBSA10
CBSA6/7_7/9
CBSA7/9_10
Triple-KO
D
Deletion stabilty over time
Deletion abundance (>10bp)
Day 2
Day 7
Day 14
CBSA6/7
CBSA7/9
CBSA10

**Figure S4**

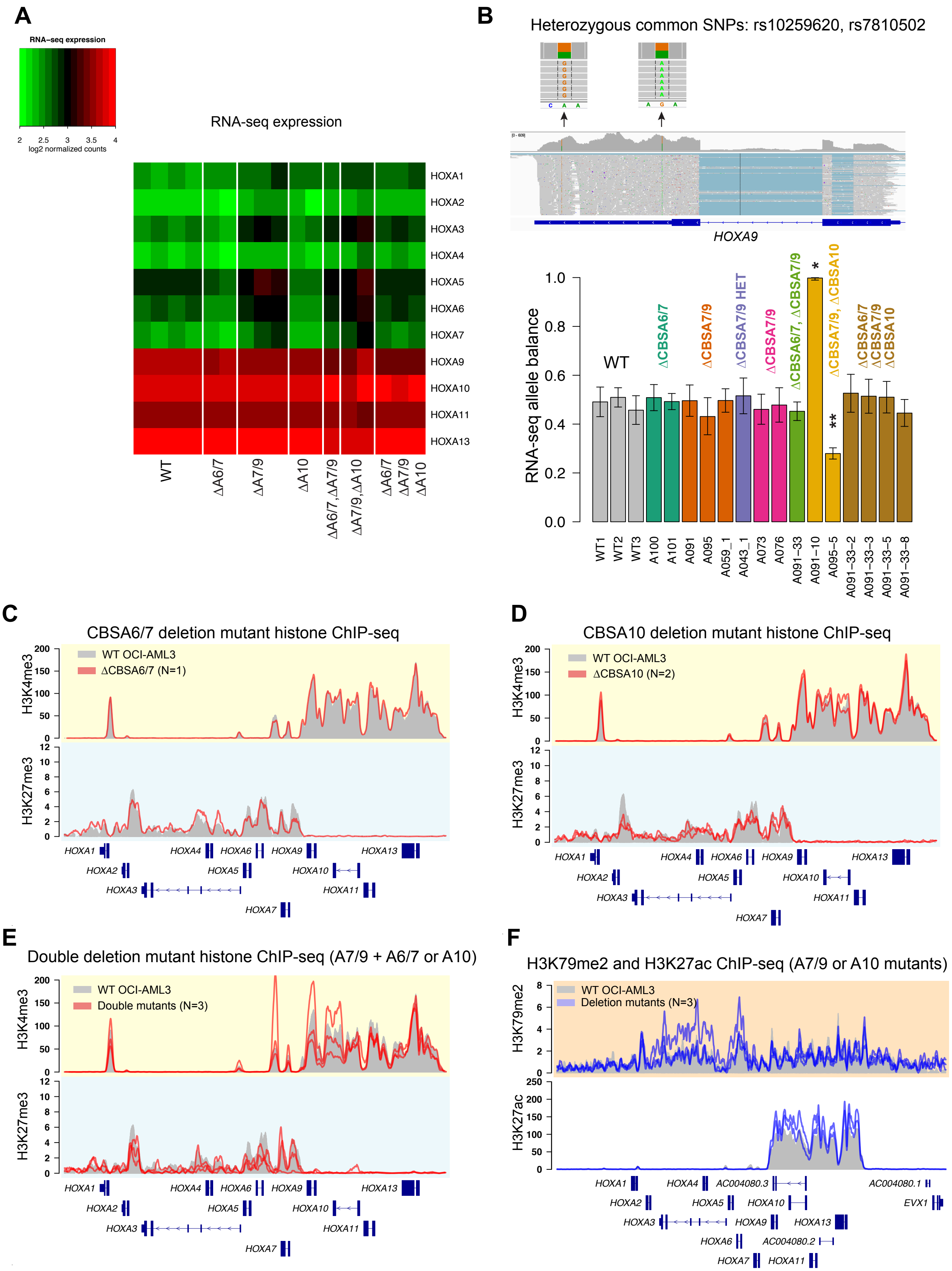
A
RNA-seq expression
log2 normalized counts
RNA-seq expression
HOXA1
HOXA2
HOXA3
HOXA4
HOXA5
HOXA6
HOXA7
HOXA9
HOXA10
HOXA11
HOXA13
WT
ΔA6/7
ΔA7/9
ΔA10
ΔA6/7,ΔA7/9
ΔA7/9,ΔA10
ΔA6/7 ΔA7/9 ΔA10
B
Heterozygous common SNPs: rs10259620, rs7810502
HOXA9
RNA-seq allele balance
WT
ΔCBSA6/7
ΔCBSA7/9
ΔCBSA7/9 HET
ΔCBSA7/9
ΔCBSA6/7, ΔCBSA7/9
ΔCBSA7/9, ΔCBSA10
ΔCBSA6/7 ΔCBSA7/9 ΔCBSA10
WT1
WT2
WT3
A100
A101
A091
A095
A059_1
A043_1
A073
A076
A091−33
A091−10
A095−5
A091−33−2
A091−33−3
A091−33−5
A091−33−8
C
CBSA6/7 deletion mutant histone ChIP-seq
WT OCI-AML3
ΔCBSA6/7 (N=1)
H3K4me3
H3K27me3
D
CBSA10 deletion mutant histone ChIP-seq
WT OCI-AML3
ΔCBSA10 (N=2)
E
Double deletion mutant histone ChIP-seq (A7/9 + A6/7 or A10)
WT OCI-AML3
Double mutants (N=3)
F
H3K79me2 and H3K27ac ChIP-seq (A7/9 or A10 mutants)
WT OCI-AML3
Deletion mutants (N=3)
H3K79me2
H3K27ac
AC004080.3
AC004080.1
AC004080.2
EVX1

### Figure S5

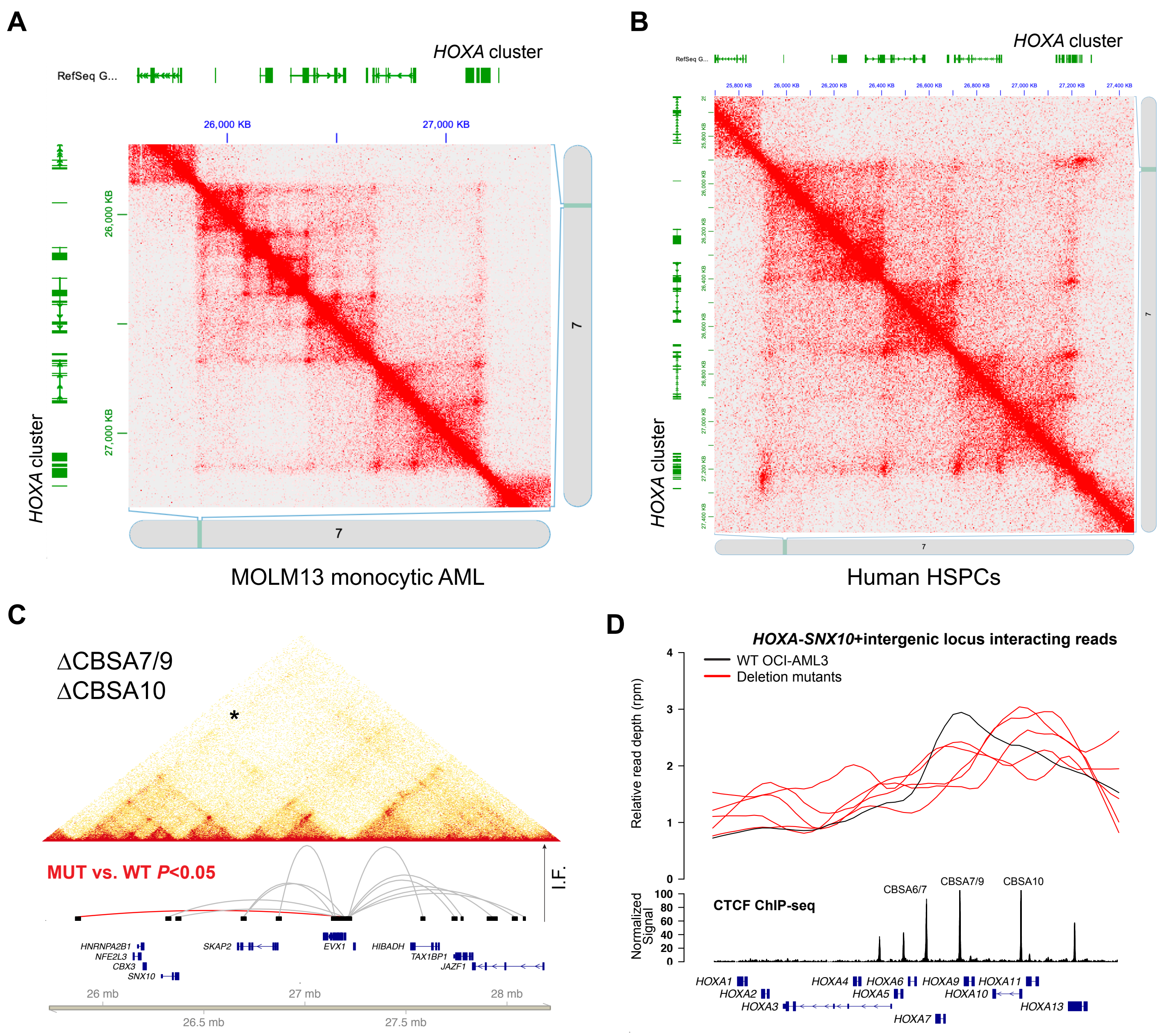
A
HOXA cluster
RefSeq G...
26,000 KB
27,000 KB
7
MOLM13 monocytic AML
B
HOXA cluster
7
Human HSPCs
C
ΔCBSA7/9
ΔCBSA10
*
MUT vs. WT P<0.05
I.F.
HNRNPA2B1
NFE2L3
CBX3
SNX10
SKAP2
EVX1
HIBADH
TAX1BP1
JAZF1
26 mb
27 mb
28 mb
26.5 mb
27.5 mb
D
HOXA-SNX10+intergenic locus interacting reads
WT OCI-AML3
Deletion mutants
Relative read depth (rpm)
CTCF ChIP-seq
Normalized Signal
CBSA6/7
CBSA7/9
CBSA10
HOXA1
HOXA2
HOXA3
HOXA4
HOXA5
HOXA6
HOXA7
HOXA9
HOXA10
HOXA11
HOXA13

**Figure S6**

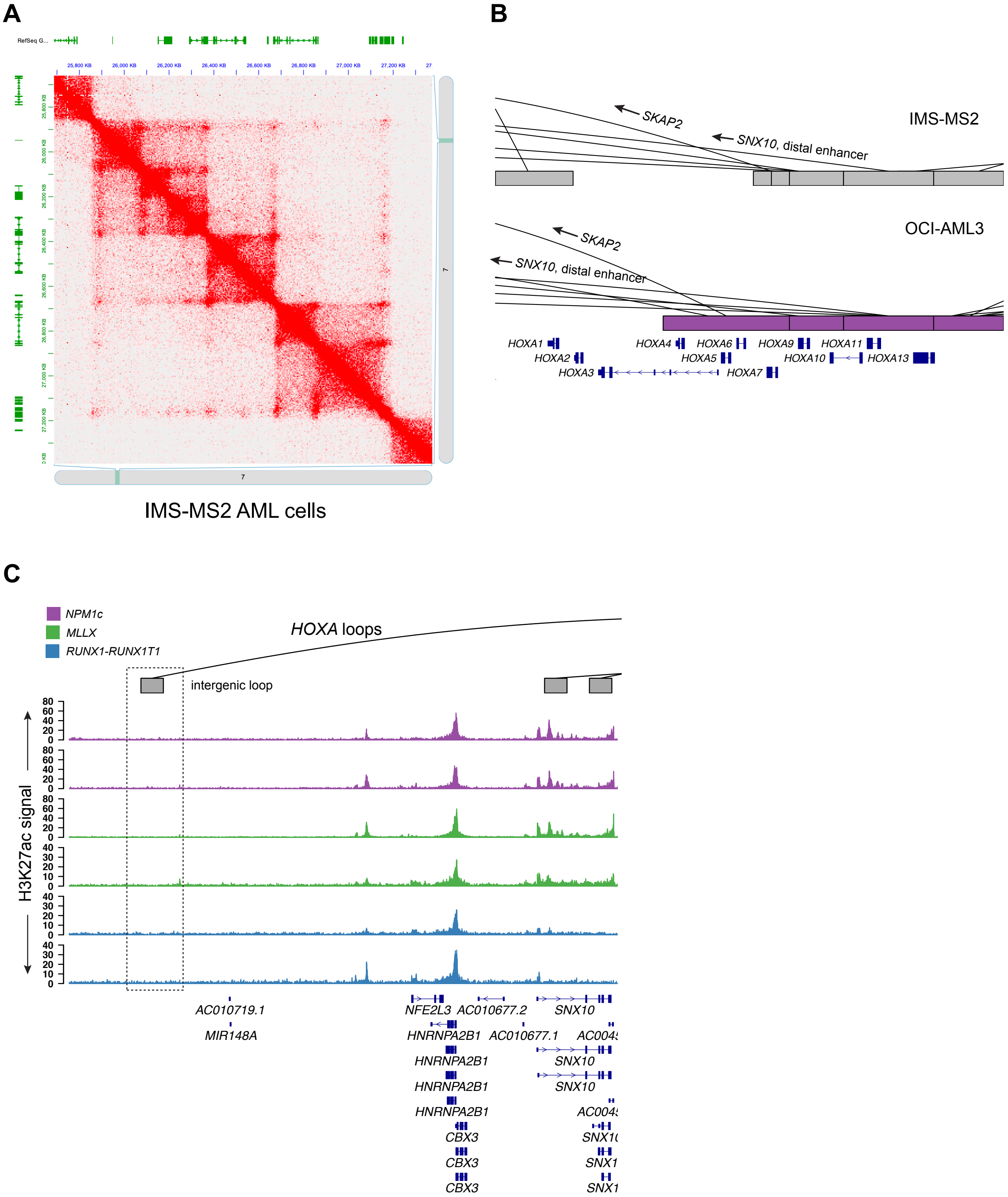
A
RefSeq G...
25,800 KB
26,000 KB
26,200 KB
26,400 KB
26,600 KB
26,800 KB
27,000 KB
27,200 KB
7
IMS-MS2 AML cells
B
SKAP2
SNX10, distal enhancer
IMS-MS2
OCI-AML3
HOXA1
HOXA2
HOXA3
HOXA4
HOXA5
HOXA6
HOXA7
HOXA9
HOXA10
HOXA11
HOXA13
C
NPM1c
MLLX
RUNX1-RUNX1T1
HOXA loops
intergenic loop
H3K27ac signal
AC010719.1
MIR148A
NFE2L3
AC010677.2
AC010677.1
HNRNPA2B1
CBX3
SNX10
